# Diagnosis of Human Leptospirosis: High Resolution Melting Analysis for Direct Detection of *Leptospira* in the Early Stage of the Disease in a Clinical Setting

**DOI:** 10.1101/311142

**Authors:** Lisa M. Esteves, Sara M. Bulhões, Claudia C. Branco, Teresa Carreira, Maria L. Vieira, Maria Gomes-Solecki, Luisa Mota-Vieira

## Abstract

Currently, the direct detection of *Leptospira* infection can be done in clinical laboratories by a conventional nested polymerase chain reaction method (nested PCR), which is labourious and time-consuming. To overcome these drawbacks, we tested a set of paired samples of serum and urine from 202 patients presenting at a hospital located in an area endemic for leptospirosis using high resolution melting (HRM). The results were compared with those obtained by nested PCR for direct detection of the pathogen in both specimens and with the gold standard test used for indirect detection of anti-*Leptospira* antibodies in serum (the microscopic agglutination test, MAT). The HRM assay results were positive for 46/202 (22.7%) samples, whereas 47/202 (23.3%) samples were positive by nested PCR. As expected in recently infected febrile patients, MAT results were positive in only 3/46 (6.5%) HRM-positive samples. We did a unique comparative analysis using a robust biobank of paired samples of serum and urine from the same patient to validate the HRM assay for molecular diagnosis of human leptospirosis in a clinical setting. This assay fills a void unmet by serologic assays as it can detect the presence of *Leptospira* in biological samples even before development of antibody takes place.

## Introduction

Leptospirosis is a worldwide zoonotic and neglected infectious disease caused by pathogenic bacteria of the *Leptospira* genus from the family Leptospiraceae^1^. This disease is known for its endemicity mainly in countries with a humid tropical or subtropical climate, such as Brazil, India and Portugal (Azores Islands)^2^. The infection is associated with a variety of clinical manifestations, ranging from flu-like symptoms to multiple organ failure and death. As a result, the disease is often difficult to diagnose clinically, and laboratory support is essential^3^. Treatment with appropriate antibiotics should be initiated as early as possible after laboratory confirmation; however, the majority of patients suspected to have leptospirosis are treated empirically with broad-spectrum antibiotics effective against most bacteria before a definitive diagnosis is established. At the Hospital of Divino Espírito Santo of Ponta Delgada (HDES), located on São Miguel Island (Azores), *Leptospira* infection is confirmed in the laboratory by identifying the presence of specific fragments of *Leptospira* DNA in patient samples (serum and urine) through conventional nested polymerase chain reaction (nested PCR)^4,5^. This method is time-consuming (it takes approximately 5 hours) and is sometimes too slow to support the clinical decision for antibiotic therapy.

Current techniques to detect *Leptospira* infection are evolving from conventional PCR to real-time PCR, which is faster, tends to have higher sensitivity and specificity at detecting pathogenic *Leptospira* species and is performed in a closed system that reduces the risk of DNA cross-contamination^6^. An emerging technique for clinical diagnosis is high resolution melting (HRM) analysis. HRM was first described by Carl Wittwer’s group for mutation screening^7^, and the underlying principle is the generation of different melting curve profiles due to sequence variations in double-stranded DNA. HRM is typically performed with a real-time PCR instrument immediately after PCR. Advantages of this method include a rapid turn-around time (less than 2hr), a closed-tube format that significantly reduces contamination risk, high sensitivity and specificity, low cost and, unlike other methods, no sample processing or separations after PCR^8^. Furthermore, HRM is a non-damaging method that enables the subsequent analysis of the sample by other methods, such as DNA sequencing or gel electrophoresis^9^.

In a clinical diagnostic context, HRM has been validated for the detection of oncogene mutations^10^, human malaria diagnosis^11^, species differentiation and genotyping within microbial species^12^, but not diagnosis of human leptospirosis. Recently, two studies described an HRM method for typing *Leptospira* strains at the species and subspecies levels^13,14^; the method described in the first study can accurately discriminate *L. interrogans*, *L. kirschneri*, *L. borgpetersenii* and *L. mayottensis* with a specificity and reproducibility of 100% and less than 0.5% variation in the melting temperature (Tm) coefficient^13^. The second study describes a PCR-HRM assay that targets the 16S ribosomal gene to identify *Leptospira* species from isolated cultures^14^. However, in both studies, HRM was not evaluated in patient samples as a clinical diagnostic test for human leptospirosis.

The aim of the present work was to evaluate a diagnostic assay for human leptospirosis capable of providing timely laboratory results on the same day the patient is seen at the emergency room. To address this unmet need, we used a robust biobank of paired serum and urine samples and evaluated the accuracy of HRM analysis as a clinical diagnostic tool for direct detection of *Leptospira* in the very early stage of human leptospirosis.

## METHODS

### Ethical considerations

The present study followed international ethical guidelines and was evaluated and approved by the Health Ethics Committee of the HDES (Ref. HDES/CES/159/2009). The analysis of retrospective samples (serum and urine) from patients suspected of having leptospirosis was exempted from the need to obtain informed consent under the regulations of the Portuguese Data Protection Commission - law 12/2005 article 19, number 6 (https://www.cnpd.pt/bin/orientacoes/DEL227-2007-ESTUDOS-CLINICOS.pdf, accessed February 22, 2017).

### Study design

The present work is a retrospective hospital-based study that includes samples from patients suspected of leptospirosis infection who mainly presented at the Emergency Department (n = 167, 82.7%) of the Hospital of Divino Espírito Santo of Ponta Delgada in São Miguel, Azores, a Portuguese island in which leptospirosis is endemic. A total of 202 patients were investigated from January 2015 to June 2016 (Supplementary Table S1). The mean patient age was 48.2 (±16.4) years. Higher rates of males (89.6%), farmers (20.3%) and unemployed persons (13.4%) were observed in the study population. Clinical diagnosis by the attending physician was based on signs and symptoms of leptospirosis, as previously described^15,16^. Briefly, physicians looked for epidemiological context, such as rural activities and direct contact with contaminated areas (rat urine), and clinical manifestations, including fever, myalgia, jaundice and coluria, before collecting biological samples (serum and urine) for molecular detection/confirmation of *Leptospira* spp. We centrifuged all sera and urine samples at 2000 rpm for 10 minutes. Bacterial DNA was automatically extracted from 400 μl of independent samples of serum (S1 and S2) and urine (U1 and U2) from each patient using the BioRobot EZ1 Advanced System (Qiagen). A total of 808 samples were processed.

### Reference molecular test (conventional nested PCR)

Conventional nested PCR was considered the reference standard for *Leptospira* spp. DNA detection in the present study. After automatic bacterial DNA extraction, the *rrs* (16S rRNA) gene was amplified as previously described^4,5^ by conventional nested PCR in a Biometra^®^ T-Gradient thermal cycler. We used two primer sets: forward-A 5’-GGCGGCGCGTCTTAAACATG-3’ and reverse-B 5’-TTCCCCCCATTGAGCAAGATT-3’ for the first PCR; nested-A 5’-TGCAAGTCAAGCGGAGTAGC-3’ and nested-B 5’-TTCTTAACTGCTGCCTCCCG-3’ for the nested PCR. The first PCR reaction contained 5 μl of bacterial DNA, 10 μM primers A and B, 100 μM dNTPs (Promega), 25 nM MgCl2 (Qiagen), 1X Q-Solution (Qiagen), 1X buffer (Qiagen), 5 U of HotStart Taq (Qiagen) and RNase-free water to a final volume of 50 μl. The PCR programme started with an enzyme activation step at 95°C for 15 minutes; proceeded with 30 cycles of 94°C for 1 minute, 63°C for 1 minute and 72°C for 1 minute; and ended with a final extension step at 72°C for 10 minutes. The nested PCR (2^nd^ round) used 5 μl of the first-round PCR product and 10 μM nested-A and nested-B primers. The first cycle consisted of denaturation at 95°C for 15 minutes, followed by 30 cycles of denaturation at 94°C for 1 minute, primer annealing at 63°C for 1 minute, and extension at 72°C for 1 minute, with an additional step at 72°C for 10 minutes at the end, resulting in a 292 bp fragment. Amplified *Leptospira* DNA was visualized in an UV transilluminator instrument (BioRad) after agarose gel electrophoresis (3%). A patient was defined as having a laboratory-confirmed case of leptospirosis when *Leptospira* DNA was detected in at least one serum (S1 or S2) or urine (U1 or U2) sample.

### High resolution melting (HRM) analysis

Primer pairs for HRM analysis were chosen according to the results obtained by Naze *et al*^13^. We used the following LFB1 F/R and G1/G2 primers to amplify the *lfb1* and *secY* genes, respectively: LFB1-F 5’-CATTCATGTTTCGAATCATTTCAAA-3’ and LFB1-R 5’-GGCCCAAGTTCCTTCTAAAAG-3’, and G1 5′-CTGAATCGCTGTATAAAAGT-3’ and G2 5′-GGAAAACAAATGGTCGGAAG-3’. The 15 μl reactions contained 7.5 μl of 2X Type-it HRM master mix (Qiagen), 0.7 μM final concentration of each primer (TibMolBiol), 3.75 μl of extracted bacterial DNA, and RNase-free water to a final volume of 15 μl. We performed the following amplification protocol in the 7500 Fast Real-Time PCR instrument (Applied Biosystems): denaturation at 95°C for 5 minutes, followed by 45 cycles of 95°C for 10 seconds, 55°C for 30 seconds, and 72°C for 10 seconds. These conditions were used for both primer sets. After PCR cycling, the samples were heated from 70°C to 95°C with continuous data acquisition.

We used six pathogenic *Leptospira* reference cultures provided by the Portuguese Reference Laboratory for Leptospirosis (at the Instituto de Higiene e Medicina Tropical, IHMT, of the Universidade Nova de Lisboa) as positive controls: 4 strains belonging to *L. interrogans* serogroup (sg) Icterohaemorrhagiae, *L. borgpetersenii* sg Ballum, *L. kirschneri* sg Cynopteri and *L. noguchii* sg Panama and 2 human Azorean isolates^4^ belonging to *L. interrogans* serovar (sv) Copenhageni of Icterohaemorrhagiae sg (human isolate 1) and *L. borgpetersenii* sv Arborea of Ballum sg (human isolate 6). Melting curve plots were generated and analysed using High Resolution Melt software v3.0.1 (Applied Biosystems) to determine average melting temperature (Tm) for each *Leptospira* spp.

### HRM benchmarking confirmation by Sanger sequencing

To validate the HRM analysis, we selected 18 biological specimens (13 serum and 5 urine samples) from laboratory-confirmed leptospirosis patients, including the sample positive by nested PCR and negative by HRM analysis. As reference DNA sequences, we used two *Leptospira* spp. *(L. interrogans* sg Icterohaemorrhagiae and *L. borgpetersenii* sg Ballum) and two human isolates. Amplified DNA products of *Leptospira* obtained by nested PCR were purified using the QIAquick PCR Purification Kit (Qiagen) according to the manufacturer’s instructions. Sequencing was performed using the nested-A and nested-B primer pair with the BigDye Terminator v1.1 cycle sequencing kit (Applied Biosystems) under the following conditions: 2 μl of ready reaction mix, 4 μl of BigDye sequencing buffer, 3.2 pmol of each primer pair, 7 ng of DNA, and RNase-free water to a final reaction volume of 20 μl. The cycling programme included an initial denaturation step at 96°C for 1 minute, followed by 25 cycles of 96°C for 10 seconds, 50°C for 5 seconds and 60°C for 4 minutes, in a GeneAmp^®^ PCR System 2700 (Applied Biosystems). The sequencing products were purified with a BigDye XTerminator^®^ Purification Kit (Applied Biosystems) and separated by capillary electrophoresis in an automated sequencer (ABI 3130 Genetic Analyzer, Applied Biosystems) with a 36 cm capillary and POP-7™ polymer according to the manufacturer’s instructions. Data were analysed with Sequencing Analysis software v5.3.1 (Applied Biosystems). Sequences were aligned using Bioedit™ software v7.0.0.

### Microscopic agglutination test (MAT)

A total of 46 serum samples evaluated as positive by the molecular approach were aliquoted and stored at −20°C for further detection of anti-*Leptospira* spp. antibodies by MAT. Additionally, 20 negative serum samples were selected as controls. MAT was performed at the Portuguese Reference Laboratory for Leptospirosis (IHMT, Universidade Nova de Lisboa) using a battery of 25 live pathogenic serovars (including 4 Azorean isolates) representative of 15 serogroups of pathogenic *Leptospira* and a saprophytic serovar of *L. biflexa* as an internal control. Samples were initially screened at a 1:40 dilution, and reactive sera were further diluted in a 2-fold series to the endpoint, defined as the highest serum dilution that agglutinated at least 50% of leptospires. For the Azorean endemic region, samples were considered positive when titres were 1:160 or greater, not conclusive when titres were below 1:160 (cut-off), and negative when no agglutination was observed.

### Statistical analysis

The nested PCR test was used as the reference molecular test to calculate the sensitivity, specificity, positive and negative predictive values (PPV and NPV), and overall accuracy [with the 95% confidence interval (CI)]. Calculations were performed using Vassar College’s VassarStats Website for Statistical Computation (http://www.vassarstats.net, last accessed November 10, 2017). To determine whether there was a significant difference between the diagnostic tests for *Leptospira* detection, data were analysed by McNemar’s test, and *p* < 0.05 indicated statistical significance. The Standards for Reporting of Diagnostic Accuracy (STARD) statement was followed when reporting the results of the present study^17^.

## Results

### HRM assay

The HRM assay was able to successfully distinguish 4 *Leptospira* spp. *(L. interrogans* sg Icterohaemorrhagiae, *L. borgpetersenii* sg Ballum, L. *kirschneri* sg Cynopteri and *L. noguchii* sg Panama) and the 2 human *Leptospira* isolates (HI1 and HI6). As shown in the derivative plot (Fig. 1), the LFB1 F/R and G1/G2 primer sets produced distinct melting curve profiles for reference *Leptospira* strains of *L. interrogans* and *L. borgpetersenii* spp. that matched those of the human *Leptospira* isolates (HI1 and HI6) of the same species. The Tm values obtained for LFB1 F/R were 80.71°C (*L. interrogans*), 81.84°C (*L. noguchii*), 82.31°C (*L. kirschneri*) and 83.26°C (*L. borgpetersenii*), and those for G1/G2 were 78.61°C (*L. noguchii*), 79.10°C (*L. interrogans*), 79.19°C (*L. kirschneri*) and 81.50°C (*L. borgpetersenii*). Moreover, these results were reproducible across 10 independent melt curve runs.

**Figure 1.**
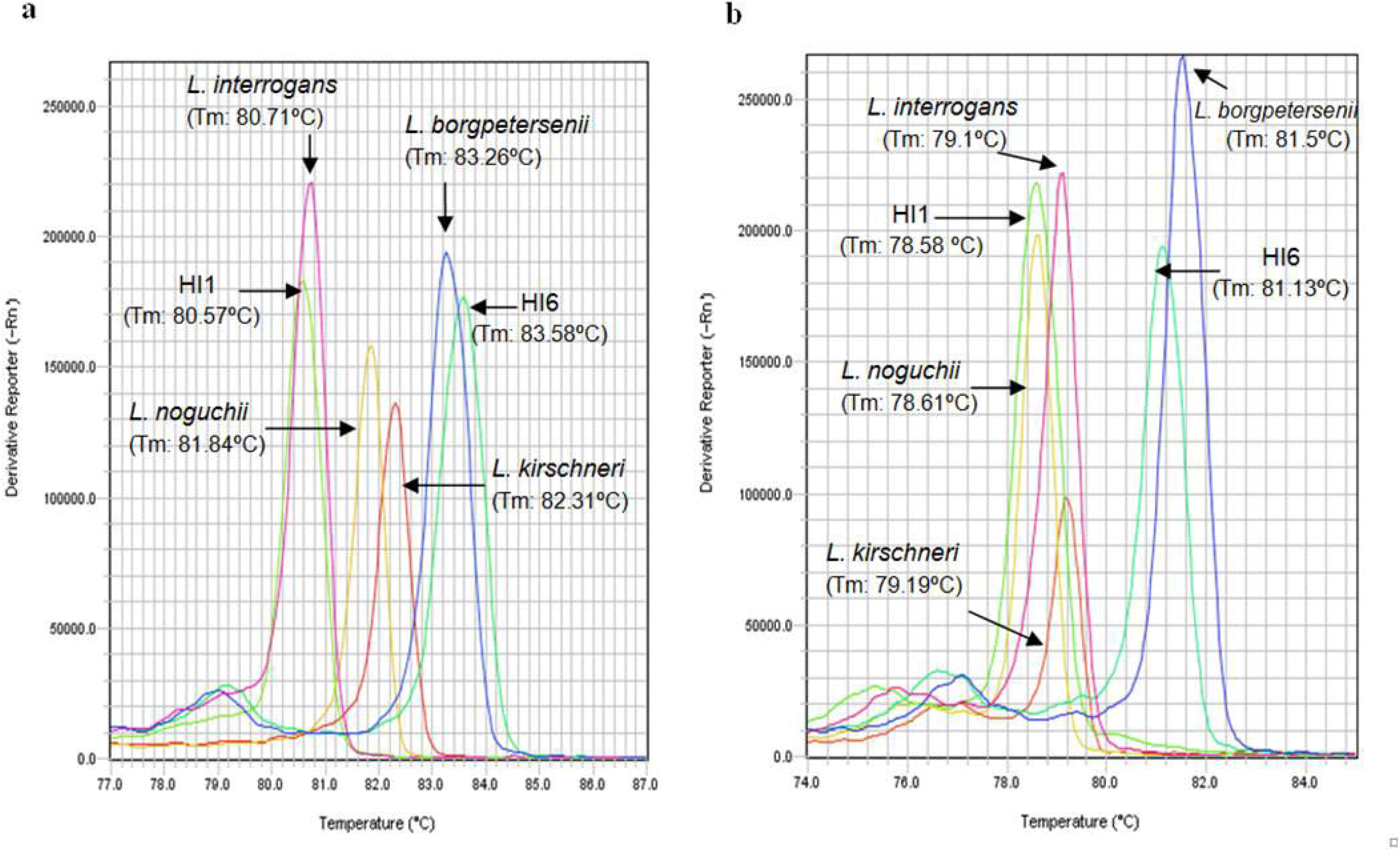
High resolution melting curve analysis profiles of cultured *Leptospira* spp., and *Leptospira* isolates from human leptospirosis patients. (**a**) HRM profiles using the LFB1 primer pair; (**b**) HRM profiles using the G1/G2 primer pair. Abbreviations: HI1, Human isolate 1; HI6, Human isolate 6.

### HRM screening of samples from patients suspected of having leptospirosis

We screened 808 clinical specimens (404 serum and 404 urine; paired samples from 202 patients) using HRM analysis. The average Tm with the LFB1 F/R primers was 80.94°C in 28 (60.9%) and 83.84°C in 16 (39.1%) patients (Supplementary Table S2). The average Tm with the G1/G2 primers was 79.36°C in 28 (60.9%) and 81.82°C in 18 (39.1%) patients. For both primer sets, we found one clinical sample to be positive by nested PCR and negative by HRM. The Tm obtained with *Leptospira* spp. and the melting curve profile results were consistent for the remaining patient samples (Fig. 2). We clustered the melting curves in two groups and identified the *Leptospira* spp. in the patient samples by comparing the Tm values to those of the six *Leptospira* positive controls.

**Figure 2.**
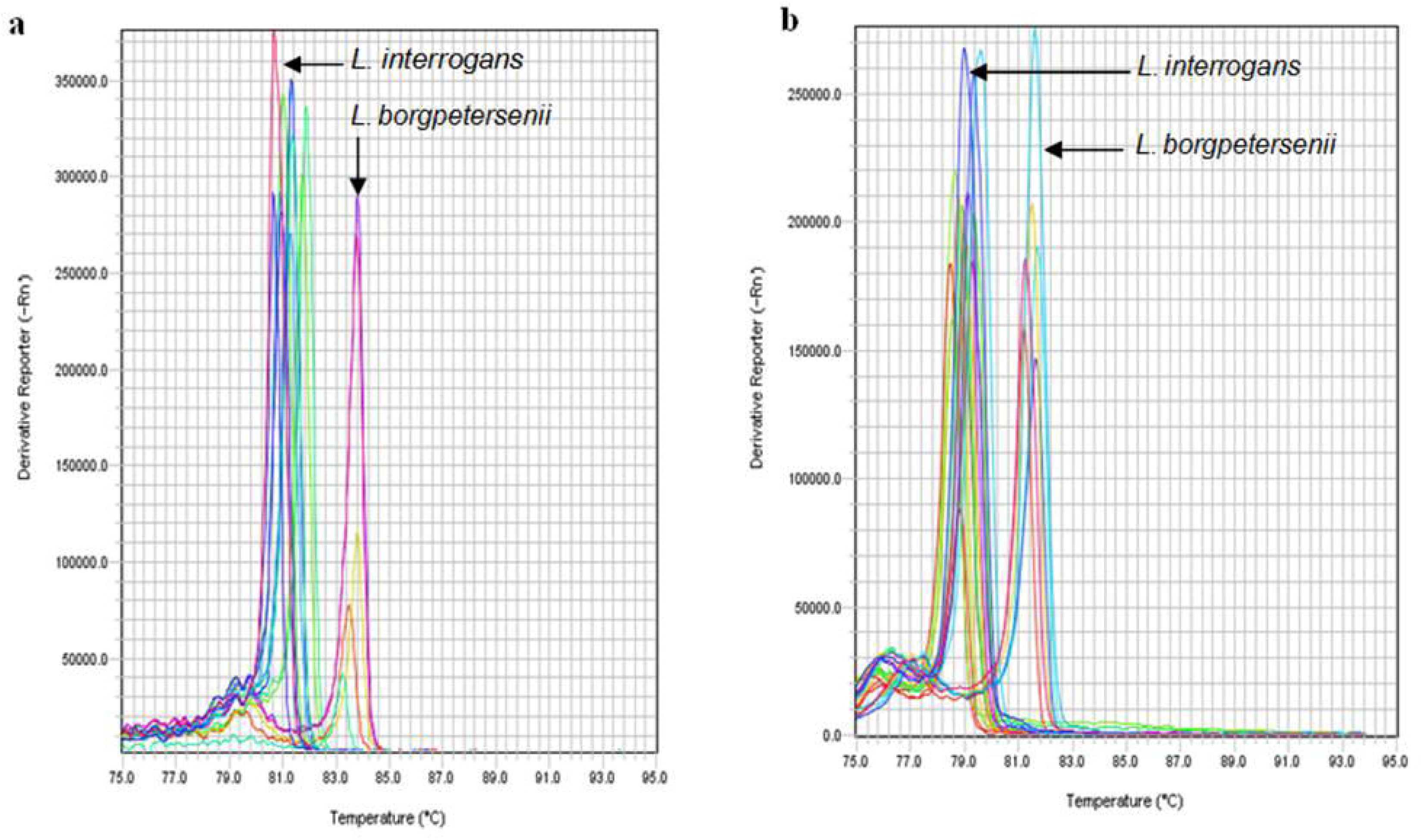
High resolution melting curve analysis profiles of human clinical samples (serum and urine) from patients with suspected leptospirosis. (**a**) HRM profiles using the LFB1 primer pair; (**b**) HRM profiles using the G1/G2 primer pair.

### Kinetics of disease progression based on *Leptospira* detection

We evaluated paired serum and urine samples from 202 patients clinically suspected of having *Leptospira* spp. infection by nested PCR and HRM (Table 1). The nested PCR results were positive for 23.3% (47/202) patients and negative for 76.7% (155/202). Using HRM, the results were positive for 22.7% (46/202) patients and negative for 77.2% (156/202). The discrepant result between the two molecular assays was confirmed to be a false positive by sequencing (see below). HRM produced conclusive results in about half of the time (~2hr) needed to generate nested PCR results (usually ~5hr).

**Table 1.**
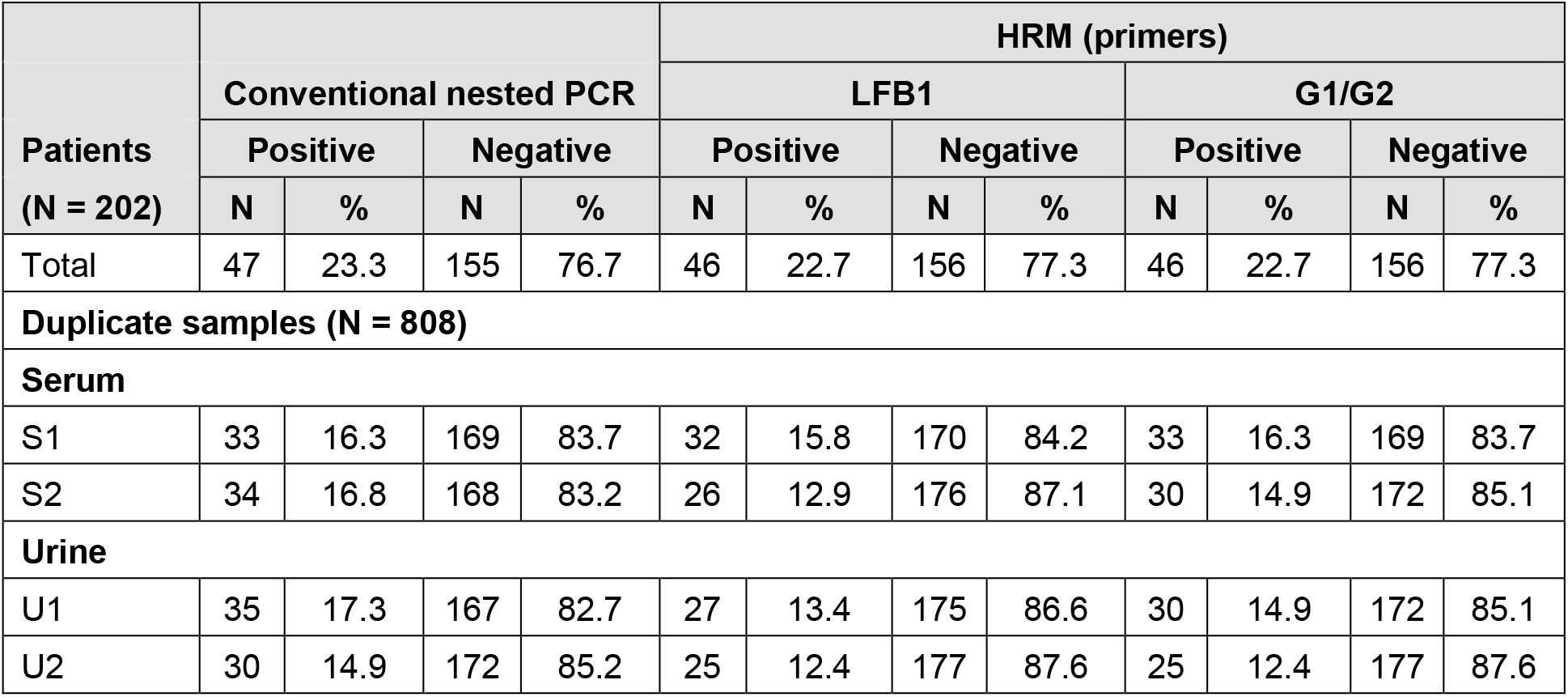
Molecular characterization of 202 patients with suspected clinical leptospirosis. Duplicate serum and urine samples were investigated by conventional nested PCR and HRM methods.

| Patients<br>(N = 202) | Conventional nested PCR |  |  |  | HRM (primers) |  |  |  |  |  |  |  |
| --- | --- | --- | --- | --- | --- | --- | --- | --- | --- | --- | --- | --- |
|  |  |  |  |  | LFB1 |  |  |  | G1/G2 |  |  |  |
|  | Positive |  | Negative |  | Positive |  | Negative |  | Positive |  | Negative |  |
|  | N | % | N | % | N | % | N | % | N | % | N | % |
| Total | 47 | 23.3 | 155 | 76.7 | 46 | 22.7 | 156 | 77.3 | 46 | 22.7 | 156 | 77.3 |
| **Duplicate samples (N = 808)** |  |  |  |  |  |  |  |  |  |  |  |  |
| **Serum** |  |  |  |  |  |  |  |  |  |  |  |  |
| S1 | 33 | 16.3 | 169 | 83.7 | 32 | 15.8 | 170 | 84.2 | 33 | 16.3 | 169 | 83.7 |
| S2 | 34 | 16.8 | 168 | 83.2 | 26 | 12.9 | 176 | 87.1 | 30 | 14.9 | 172 | 85.1 |
| **Urine** |  |  |  |  |  |  |  |  |  |  |  |  |
| U1 | 35 | 17.3 | 167 | 82.7 | 27 | 13.4 | 175 | 86.6 | 30 | 14.9 | 172 | 85.1 |
| U2 | 30 | 14.9 | 172 | 85.2 | 25 | 12.4 | 177 | 87.6 | 25 | 12.4 | 177 | 87.6 |

Based on the results of the laboratory-confirmed leptospirosis cases (n = 46), we established the kinetic profiles of disease progression (Table 2). Profile A characterizes patients who had a positive molecular result in serum and a negative result in urine, which represents early dissemination of *Leptospira* in blood. In Profile B patients were positive in both serum and urine, which represents the transit of *Leptospira* infection to the kidney and other tissues. Profile C corresponds to patients who were positive in urine and negative in serum, which represents the *Leptospira* excretion stage. The highest percentage of patients analysed by nested PCR (60.9%) and HRM (47.8%) fell under profile B, positive for both serum and urine. The profile analysis also revealed differences between HRM and MAT, which is in accordance with the kinetics of leptospirosis progression.

**Table 2.**
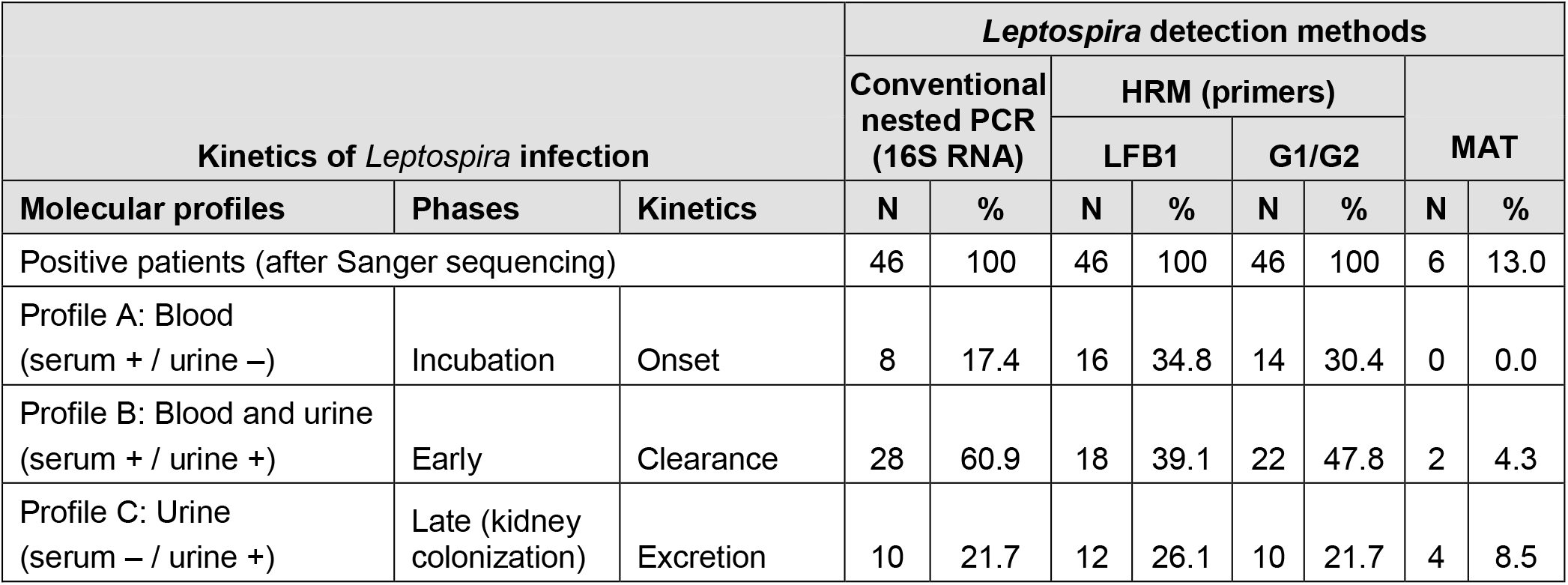
**Molecular profiles of patients with laboratory-confirmed leptospirosis and the corresponding** disease **kinetics**. *P* > 0.05, conventional nested PCR compared with HRM; *P* < 0.0001, HRM compared with MAT.

### Sequencing analysis

To benchmark *Leptospira* detection by HRM analysis, we performed Sanger sequencing (Fig. 3). The obtained bacterial DNA sequences confirmed the positive HRM results in 17 clinical samples. One sample (#18) was positive in the urine by nested PCR but negative by HRM. This sample was assessed twice by sequencing, HRM and nested PCR, and the sequence had a 97% match to *Collinsella aerofaciens*, which is found predominantly in the human gut. For the remaining samples (17/18), we observed a perfect match with the reference sequences regarding Tm values, melting curve profiles, and sequencing data.

**Figure 3.**
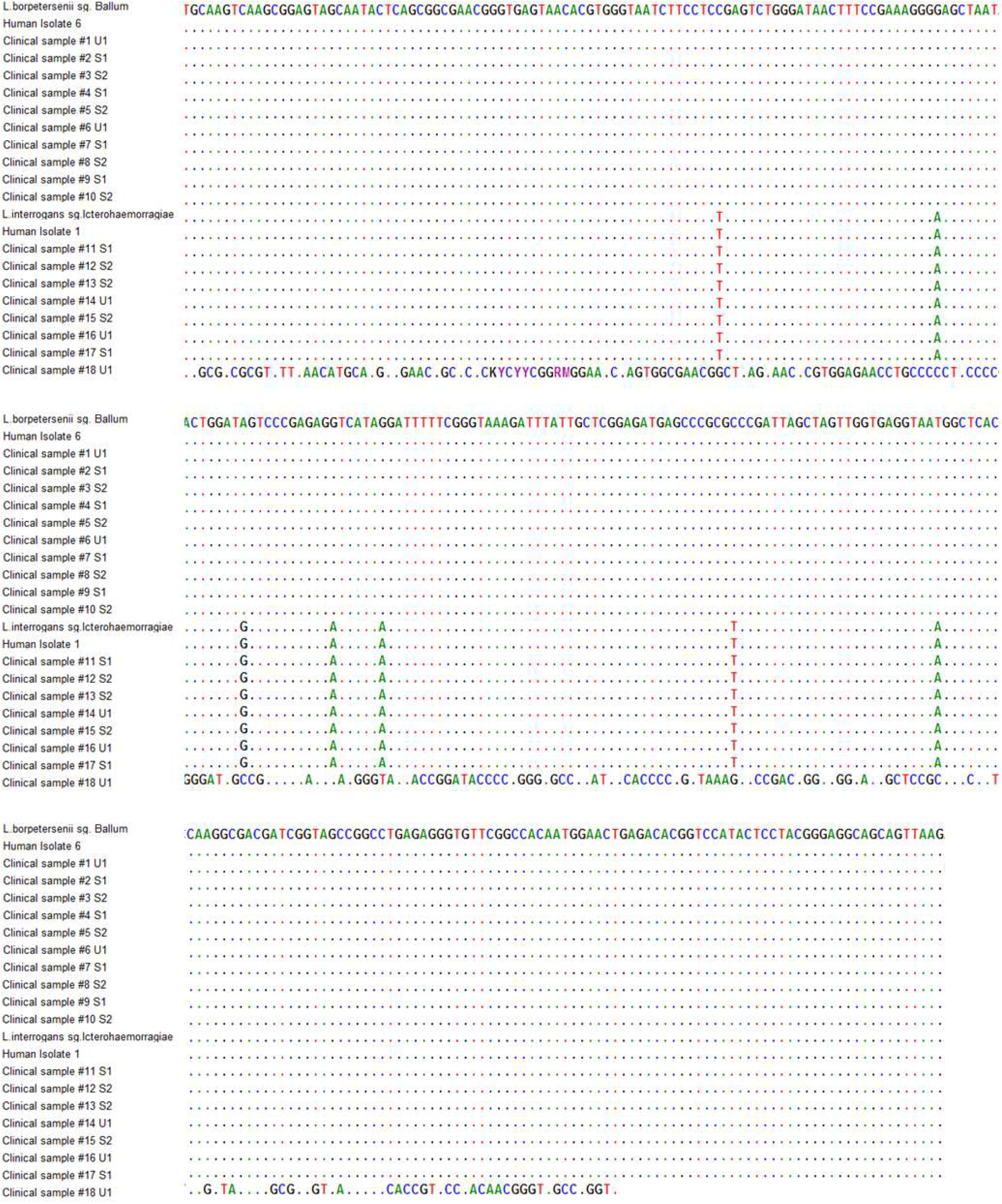
Confirmation of the HRM analysis by Sanger sequencing. Alignment of the consensus sequences of the clinical samples, *Leptospira* spp. and human isolates. Only the sequences showing differences from the first sequence are shown. Nucleotides identical to those in the first sequence are indicated by dots.

### Microscopic agglutination test (MAT)

MAT results revealed that of the 46 nested PCR-positive patients, only 3 presented specific *anti-Leptospira* antibodies (6.5%), and 3 presented *anti-Leptospira* antibodies below the cut-off titre adopted by the Portuguese Reference Laboratory for Leptospirosis in the Azorean endemic region (Table 3).

**Table 3.** MAT results from the 46 patients with laboratory-confirmed leptospirosis. Abbreviations: Arb A [serovar Arborea (Azorean isolate) serogroup Ballum]; Cyn [Cynopteri (reference serovar) serogroup Cynopteri]; NC, not conclusive (specific reactivity below the cut-off of 1:160 adopted in the Azorean endemic region).

| Kinetics of *Leptospira* infection |  | Serovar | Species |
| --- | --- | --- | --- |
| Profiles | Conventional nested PCR (N = 46) | MAT | HRM |
| C | 1 | Positive (co-agglutination – highest titre 1:1280 against Arb A and Cyn) | *L. borgpetersenii* |
|  | 1 | Positive (Arb A 1:160) | *L. borgpetersenii* |
|  | 1 | Positive (co-agglutination – highest titre 1:320 against Arb A 1:320) | *L. interrogans* |
|  | 1 | NC | *L. borgpetersenii* |
|  | 6 | Negative | *L. interrogans* |
| B | 1 | NC | *L. interrogans* |
|  | 1 | NC | *L. interrogans* |
|  | 16 | Negative | *L. interrogans* |
|  | 10 | Negative | *L. borgpetersenii* |
| A | 3 | Negative | *L. interrogans* |
|  | 5 | Negative | *L. borgpetersenii* |

### Analytical specificity and sensitivity of HRM

To validate the HRM analysis as a diagnostic method for *Leptospira* spp. detection, we assessed the accuracy parameters by comparing the results of nested PCR (reference molecular test) after sequencing with those obtained by HRM (Table 4). Of the 46 patients who were positive for leptospirosis by nested PCR, 46 had a positive HRM result, for a sensitivity of 1.00 (95% CI: 0.90-1.00). Moreover, of the 156 patients who were negative for leptospirosis by nested PCR, 156 had a negative HRM result, for a specificity of 1.00 (95% CI: 0.97-1.00). The PPV and NPV were 1.00 (95% CI: 0.90-1.00) and 1.00 (95% CI: 0.97-1.00), respectively. Overall, HRM accuracy was 100%. Together, these results confirm and validate the accuracy of HRM as a clinical diagnostic test for human leptospirosis.

**Table 4.** Diagnostic accuracy of HRM analysis compared with conventional nested PCR for detecting human leptospirosis infection. Abbreviation: CI, confidence interval.

| Patients (N = 202) | HRM |  |
| --- | --- | --- |
|  | Estimated value | 95% CI |
| Sensitivity | 1.00 | 0.90–1.00 |
| Specificity | 1.00 | 0.97–1.00 |
| Positive predictive value (PPV) | 1.00 | 0.90–1.00 |
| Negative predictive value (NPV) | 1.00 | 0.97–1.00 |
| Accuracy (%) | 100 | – |

### Comparison between HRM and other molecular PCR and serological diagnostic assays for leptospirosis

The current study is the first one to present 100% accuracy values - specificity, sensitivity, Positive Predictive Values (PPV) and Negative Predictive Values (NPV) - for a molecular PCR method, validating the HRM as the best test to be implemented in a clinical setting. No other molecular test has provided PPVs and NPVs (Table 4). Compared with serological methods, HRM has the highest diagnostic value as it can be used to detect *Leptospira* directly in biological samples collected in the first days of the infection, making this test the most reliable to inform treatment decisions for hospitalized patients and patients seen in emergency rooms or clinics (Table 5).

**Table 5.**
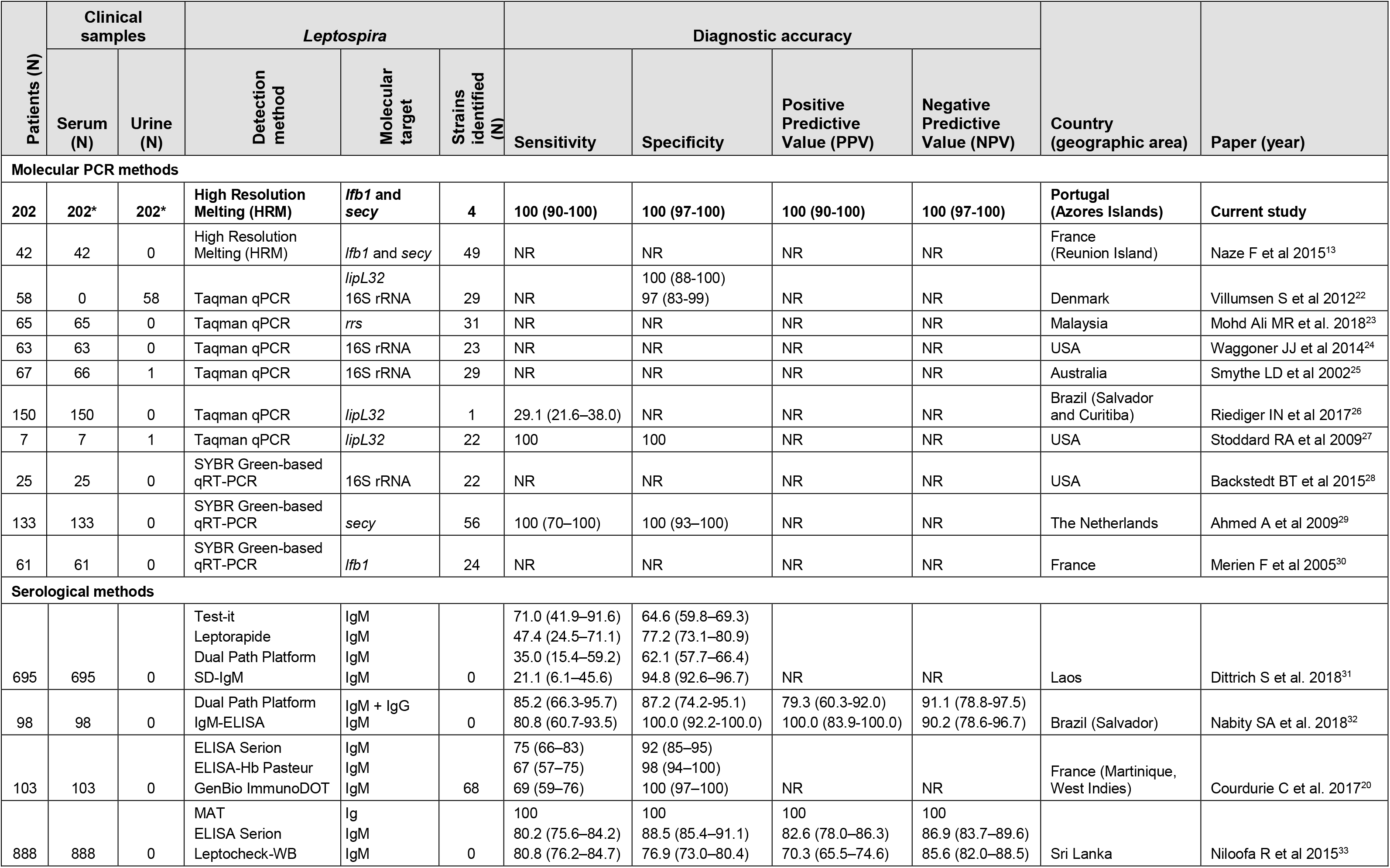
Comparison between molecular and serological assays for the diagnosis of leptospirosis. *Analysed in duplicated; NR, not reported.

## Discussion

In this study, the HRM assay was validated for the accurate detection of *Leptospira* DNA in biological samples from patients presenting in the emergency room of a hospital in the Azorean island of São Miguel, a Portuguese region endemic for leptospirosis. Among 202 human patients suspected of having leptospirosis, 46 tested positive (22.7%) by both nested PCR and HRM; among these patients, only 3 tested positive (6.5%) by MAT. Melting curve profiles with the LFB1 F/R primer set distinguished the 4 *Leptospira* spp., *L. interrogans, L. borgpetersenii, L. kirschneri* and *L. noguchii*, in cultured bacteria and human isolates (Fig. 1). These results are in accordance with those obtained by Naze and collegues^13^. In addition, template-independent amplifications targeting the two relevant genes (*lfb1* and *secY*) of pathogenic *Leptospira* spp. also provided a thorough validation of the present HRM assay. The 404 human samples used - paired serum and urine from 202 patients, analysed in duplicated (total of 808 specimens) - validate, for the first time, the application of HRM as a clinical diagnostic test for human leptospirosis in a clinical setting.

The melting curve analysis of *Leptospira* species in patient samples (serum and urine) accurately discriminated species when positive controls were included in each run (Fig. 2). According to the Tm, the HRM assay revealed that 60.9% (28/46) of patients were infected with leptospires belonging to *L. interrogans*, and 39.1% (18/46) were infected with leptospires belonging to *L. borgpetersenii* species (Supplementary Table S2). The most likely explanation for these results is that *L. interrogans* survives longer when exposed to the environment, which is why it is more prone than *L. borgpetersenii* to infect humans. The latter does not survive in the environment and is transmitted by direct contact with the host^18^. MAT is the hundred-year old gold standard method for the serodiagnosis of leptospirosis and allows for the determination of the presumptive serogroup or serovar of the infecting strain in routine diagnostics and/or epidemiological studies^19^. In the present study, MAT results substantiated the HRM findings, as these patients presented anti-*Leptospira* antibodies belonging to one of these serogroups. In addition, MAT results were positive in only 3/46 (6.5%) of the HRM-positive samples which is expected in recently infected febrile patients and explained by the typical delay period between time of infection and presence of measurable levels of antibodies in blood. Low MAT sensitivity in an early stage of disease infection was discussed previously^20^. In a clinical diagnostic context, this observation alone qualifies HRM as a valuable alternative to MAT by providing early unambiguous diagnosis of the disease, which can better inform treatment decisions by the physician as recommended by WHO^3^. The HRM method validated in the present study not only detects *Leptospira* in human biological samples with 100% accuracy, but also informs epidemiology of the disease by identifying the infecting species.

By conducting DNA sequencing as part of the assay validation, we determined that the leptospires infecting these patients belonged to the serogroups Icterohaemorrhagiae and Ballum (Fig. 3). These results agree with prior studies performed in the Azores Islands (São Miguel and Terceira), where the serogroups Icterohaemorrhagiae (*L. interrogans*) and Ballum (*L. borgpetersenii*) were the most frequent human^4,15^ and rodent *Leptospira* isolates^16,21^.

The profiles based on the 46 confirmed positive patients (by nested PCR and HRM) described in Table 2 are in accordance with the kinetics of *Leptospira* infection and disease progression in humans^18^. The analysis allows us to identify the illness point at which patients presented at the hospital. Infection produces leptospiraemia within the first days after exposure, which is followed by the appearance of leptospires in multiple organs by the 3^rd^ day of infection (incubation period and dissemination). Illness develops with the appearance of agglutinating antibodies 5-14 days after exposure (early phase). Leptospires are cleared from the bloodstream and organs in the late phase, as serum agglutinating antibody titres increase^18^. A higher percentage of patients in this study were seen in the early phase of the disease (profile B, Table 2), when the immune system starts to produce antibodies and clearing *Leptospira* from the blood, which is why the bacteria is detected in serum and urine. Another important finding is that HRM is more sensitive than nested PCR at detecting *Leptospira* during the incubation period (first seven days, profile A). This finding is of clinical relevance because it allows for the immediate initiation of antibiotic therapy at the earliest onset. Regarding profile C, 23.4% of patients presented at the hospital when *Leptospira* DNA is detected in the urine. This delay in coming to the hospital probably occurs because the symptoms are similar to those of flu, and patients stay at home and take conventional over-the-counter medicine. For patients with profile C, HRM was more specific than nested PCR; one patient was positive by nested PCR and negative by HRM. Bacterial DNA sequencing of this patient’s urine sample (#18) showed a 97% match to *Collinsella aerofaciens*, a type of bacteria found in the human gut, proving that the nested PCR result was a false positive. This finding highlights the caution necessary when interpreting the results of assays such as nested PCR that target the *rrs* gene (encoding 16S rRNA), which is conserved among many bacterial species, and are thus prone to cross-reactivity. Extra care should be taken when validating PCR assays based on the *rss* gene, especially in urine samples that contain poorly characterized microbial flora, which is supported by previous observations^22^. The performance of the HRM assay was evaluated and compared with that of the reference molecular test, nested PCR (Table 4); HRM was 100% accurate. The high specificity (100%) and sensitivity (100%) of HRM in endemic regions, such as the Azores, is highly relevant. Notably, since the nested PCR technique was implemented at HDES in 2005, no patient on São Miguel Island has died of leptospirosis. According to official data in the Azores (the islands of São Miguel and Terceira) for the period between 1986 and 2002, fewer than 19 deaths due to leptospirosis were reported each year^16^.

In clinical diagnostic laboratories, real-time PCR methods are increasingly being used instead of conventional PCR methods, providing the opportunity to rapidly confirm leptospirosis infection in the first days of infection. So far, this is the most comprehensive study performed for laboratorial diagnosis of human leptospirosis using paired samples of serum and urine from the same patient. HRM allows for accurate clinical diagnosis of leptospirosis in just 2 hours, rather than the 5 hours needed for nested PCR, and the results are unambiguous and easy to interpret. The HRM assay is a robust molecular PCR method for the diagnosis of human leptospirosis infection in endemic regions and it can be fully implemented in routine clinical laboratories with real-time PCR equipment. Furthermore, HRM has the advantage of allowing for the distinction of *Leptospira* species which informs leptospirosis epidemiology in the geographic region without requiring the maintenance of large strain collections and labourious cultures. Recently, molecular PCR and serological methods for the diagnosis of human leptospirosis have been published^13, 20, 23–33^. However, their accuracy values are still below those reported here. An important limitation of these serological assays is the inability to detect *anti-Leptospira* antibodies in the very early stages of infection.

In conclusion, we did a unique comparative analysis using a robust biobank of paired samples of serum and urine from the same patient to validate the HRM assay for molecular diagnosis of human leptospirosis in a clinical setting. As a clinical diagnostic method, it is imperative to use both primer sets in each run to amplify the *lfb1* and *secY* genes and to include at least one positive and one negative control. Furthermore, rapidly distinguishing *Leptospira* species while performing the diagnostic test adds an epidemiological advantage to the assay over current clinical molecular diagnostic techniques.

## Acknowledgements

The present work was supported by grants from the Direção Regional da Ciência e Tecnologia (M121/I/OLD/2002, from the Azores Government) and BioISI (Centre Reference: UID/MULTI/04046/2013) from FCT/MCTES/PIDDAC, Portugal. The authors would like to thank Rúben Neves, a Ph.D. student at Clark University, for revising the English language of the manuscript.

## Author Contributions

L.M.E. and L.M.V. designed the study and wrote the manuscript, L.M.V. provided materials and reagents, M.L.V. provided the *Leptospira* strains and human isolates, M.L.V. and T.C. performed and analysed the MAT experiments, L.M.E. performed and analysed the HRM experiments, S.M.B. performed and analysed the sequencing experiments, C.C.B. tested and calibrated High Resolution Melt software v3.0.1 (Applied Biosystems), and M.G.S. provided critical reading and editing of the manuscript. All authors read, edited and approved the final manuscript.

## Data availability

All data generated or analysed during this study are included in this published article and the Supplementary Information files.

## Additional Information

### Supplementary information

accompanies this paper.

## References

1. Levett, P. N. Leptospirosis. Clin. Microbiol. Rev. 14, 296–326 (2001).

2. Collares-Pereira, M. et al. First epidemiological data on pathogenic leptospires isolated on the Azorean islands. Eur. J. Epidemiol. 13, 435–441 (1997).

3. World Health Organization. Human leptospirosis: guidence for diagnosis, surveillance, and control. WHO Library, ISBN 924154589 5, 1–107 (2003).

4. Gonçalves, A. T. et al. First isolation of human Leptospira strains, Azores, Portugal. Int. J. Infect. Dis. 14, e148–e153, doi: 10.1016/j.ijid.2009.12.004 (2010).

5. Mérien, F., Amouriaux, P., Perolat, P., Baranton, G. & Saint Girons, I. Polymerase chain reaction for detection of Leptospira spp. in clinical samples. J. Clin. Microbiol. 30, 2219–2224 (1992).

6. Tong, S. Y. & Giffard, P. M. Microbiological applications of high-resolution melting analysis. J. Clin. Microbiol. 50, 3418–3421, doi: 10.1128/JCM.01709-12 (2012).

7. Wittwer, C. T., Reed, G. H., Gundry, C. N., Vandersteen, J. G. & Pryor, R. J. High-resolution genotyping by amplicon melting analysis using LCGreen. Clin. Chem. 49, 853–860, doi: 10.1373/49.6.853 (2003).

8. Montgomery, J. L., Sanford, L. N. & Wittwer, C. T. High-resolution DNA melting analysis in clinical research and diagnostics. Expert Rev. Mol. Diagn. 10, 219–240, doi: 10.1586/erm.09.84 (2010).

9. Li, B. S. et al. Is high resolution melting analysis (HRMA) accurate for detection of human disease-associated mutations? A meta analysis. Plos One. 6, e28078, doi: 10.1371/journal.pone.0028078 (2011).

10. Negru, S. et al. KRAS, NRAS and BRAF mutations in Greek and Romanian patients with colorectal cancer: a cohort study. BMJ Open. 4, e004652, doi: 10.1136/bmjopen-2013-004652 (2014).

11. Chua, K. H. et al. Development of high resolution melting analysis for the diagnosis of human malaria. Sci. Rep. 5, 15671, doi: 10.1038/srep15671 (2015).

12. Andini, N. et al. Microbial typing by machine learned DNA melt signatures. Sci. Rep. 7, 42097, doi: 10.1038/srep42097 (2017).

13. Naze, F. Desvars, A., Picardeau, M., Bourhy, P. & Michault, A. Use of new high resolution melting method for genotyping pathogenic Leptospira spp. Plos One. 10, e0127430, doi: 10.1371/journal.pone.0127430 (2015).

14. Peláez Sánchez, R. G., Quintero J. Á. L., Pereira, M. M. & Agudelo-Flórez, P. High-resolution melting curve analysis of the 16S ribosomal gene to detect and identify pathogenic and saprophytic Leptospira species in colombian isolates. Am. J. Trop. Med. Hyg. 96, 1031–1038, doi: 10.4269/ajtmh.16-0312 (2017).

15. Esteves, L. et al. Human leptospirosis: seroreactivity and genetic susceptibility in the population of São Miguel Island (Azores, Portugal). Plos One. 9, e108534, doi: 10.1371/journal.pone.0108534 (2014).

16. Vieira, M. L., Gama-Simões, M. J. & Collares-Pereira, M. Human leptospirosis in Portugal: a retrospective study of eighteen years. Int. J. Infect. Dis. 10, 378–386, doi: 10.1016/j.ijid.2005.07.006 (2006).

17. STARD Group. STARD 2015: an updated list of essential items for reporting diagnostic accuracy studies. BMJ. 351, h5527, doi: 10.1136/bmj.h5527 (2015).

18. Ko, AI., Goarant C. & Picardeau M. Leptospira: the dawn of the molecular genetics era for an emerging zoonotic pathogen. Nat. Rev. Microbiol. 10, 736–747, doi: 10.1038/nrmicro2208 (2009).

19. Goris, M. G. & Hartskeerl R. A. Leptospirosis serodiagnosis by the microscopic agglutination test. Curr. Protoc. Microbiol. 32, Unit 12E 15 (2014).

20. Courdurie, C., Le Govic, Y., Bourhy, P., Alexer, D. & Pailla, K. Evaluation of different serological assays for early diagnosis of leptospirosis in Martinique (French West Indies). PLoS Negl. Trop. Dis. 11, e0005678, doi: 10.1371/journal.pntd.0005678 (2017).

21. Collares-Pereira, M., Santos-Reis M., Gonçalves, L., Vieira, M. L. & Flor, L. Epidemiology and control of leptospirosis in the Azores project. Final scientific report. USA Scientific Cooperative Agreement No. 58–401–3–F185, 2003–2008 (2008).

22. Villumsen, S. et al. Novel TaqMan PCR for detection of Leptospira species in urine and blood: pit-falls of in silico validation. J. Microbiol. Methods. 91, 184–190, doi: 10.1016/j.mimet.2012.06.009 (2012).

23. Mohd, M. R. et al. Development and validation of pan-Leptospira Taqman qPCR for the detection of Leptospira spp. in clinical specimens. Mol. Cell. Probes. S0890-8508(17)30127–5 (2018).

24. Waggoner, J.J. et al. Sensitive real-time PCR detection of pathogenic Leptospira spp. and a comparison of nucleic acid amplification methods for the diagnosis of leptospirosis. Plos One. 9, e112356, doi: 10.1371/journal.pone.0112356 (2014).

25. Smythe, L.D. et al. A quantitative PCR (TaqMan) assay for pathogenic Leptospira spp. BMC Infect. Dis. 2, 13, doi: 10.1186/1471-2334-2-13 (2002).

26. Riediger, I.N. et al. Rapid, actionable diagnosis of urban epidemic leptospirosis using a pathogenic Leptospira lipL32-based real-time PCR assay. PLoS Negl. Trop. Dis. 9, e0005940, doi: 10.1371/journal.pntd.0005940 (2017).

27. Stoddard, R.A., Gee, J.E., Wilkins, P.P., McCaustland, K. & Hoffmaster, A.R. Detection of pathogenic Leptospira spp. through TaqMan polymerase chain reaction targeting the LipL32 gene. Diagn. Microbiol. Infect. Dis. 3, 247–55, doi: 10.1016/j.diagmicrobio.2009.03.014 (2009).

28. Backstedt, B.T. et al. Efficient detection of pathogenic leptospires using 16S ribosomal RNA. PLoS One. 10, e0128913, doi: 10.1371/journal.pone.0128913 (2015).

29. Ahmed, A., Engelberts, M.F., Boer, K.R., Ahmed, N. & Hartskeerl, R.A. Development and validation of a real-time PCR for detection of pathogenic Leptospira species in clinical materials. PLoS One. 4, e7093, doi: 10.1371/journal.pone.0007093 (2009).

30. Merien, F. et al. A rapid and quantitative method for the detection of Leptospira species in human leptospirosis. FEMS Microbiol. Lett. 249, 139–47 (2005).

31. Dittrich, S. et al. A prospective hospital study to evaluate the diagnostic accuracy of rapid diagnostic tests for the early detection of leptospirosis in Laos. Am. J. Trop. Med. Hyg. [Epub ahead of print], doi: 10.4269/ajtmh.17-0702 (2018).

32. Nabity, S.A. et al. Prospective evaluation of accuracy and clinical utility of the Dual Path Platform (DPP) assay for the point-of-care diagnosis of leptospirosis in hospitalized patients. PLoS Negl. Trop. Dis. 12, e0006285, doi: 10.1371/journal.pntd.0006285 (2018).

33. Niloofa, R. et al. Diagnosis of leptospirosis: comparison between Microscopic Agglutination Test, IgM-ELISA and IgM Rapid Immunochromatography Test. PLoS One. 10, e0129236, doi: 10.1371/journal.pone.0129236 (2015).

